# MXD/MIZ1 complexes activate transcription of MYC-repressed genes

**DOI:** 10.1101/842799

**Authors:** Anton Shostak, Géza Schermann, Axel Diernfellner, Michael Brunner

## Abstract

MXD proteins are transcription repressors that antagonize the E-box dependent activation of genes by MYC. MYC together with MIZ1 acts also as a repressor of a subset of genes, including cell cycle inhibitor genes such as *p15* and *p21*. A role of MXDs in regulation of MYC-repressed genes is not known. Here we report that MXDs are functionally expressed in U2OS cells and activate transcription of *p15* and *p21*, and other MYC-repressed genes. Activation of transcription was dependent on the interaction of MXDs with MIZ1, and on an intact DNA binding domain. MIZ1-binding deficient MXD mutants interacted with MAX and were active as repressors of MYC-activated genes but failed to activate MYC-repressed genes. Mutant MXDs with reduced DNA binding affinity interacted with MAX and MIZ1 but neither repressed nor activated transcription. Overexpression of MXDs attenuated proliferation of U2OS cells predominantly via MIZ1-dependent induction of *p21*. Our data show that MXDs and MYC have a reciprocally antagonistic potential to regulate transcription of mutual target genes.

## Introduction

MYC-associated factor X (MAX) and its binding partners comprise a family of basic helix-loop-helix leucine zipper (bHLH-Zip) transcription factors, which are implicated in the regulation of cell growth, proliferation, differentiation, apoptosis and tumorgenesis (Carroll et al., 2018; Conacci-Sorrell et al., 2014; Poole and van Riggelen, 2017). Complexes of MAX with MYC and its homologs MYCN and MYCL bind to enhancer-box motifs (E-boxes) and promote expression of target genes (Conacci-Sorrell et al., 2014).

Under physiological conditions MYC is expressed in response to mitogens and promotes cell growth and proliferation (Armelin et al., 1984; Carroll et al., 2018; Hasmall et al., 1997; Lutterbach and Hann, 1994; Wang et al., 2011). Elevated expression or activation of MYC is associated with uncontrolled cellular growth and proliferation and supports the development of cancer, and MYC or its homologues are overexpressed, amplified or deregulated in many cancer types (Dang, 2012; Kalkat et al., 2017). MYC has been reported to function as a regulator of specific target genes (Kress et al., 2015; Muhar et al., 2018; Sabo et al., 2014; Walz et al., 2014) and/or as a general amplifier of transcription of active genes on a genome-wide scale (Baluapuri et al., 2019; Gerlach et al., 2017; Lin et al., 2012; Nie et al., 2012). MYC interacts with several coactivators and RNA-polymerase II (Pol II) associated factors including chromatin modifiers, transcription initiation and elongation factors that are implicated in transcription activation, but also with several complexes and assemblies involved in transcriptional repression (Baluapuri et al., 2019; Kress et al., 2015; Poole and van Riggelen, 2017).

MYC has been shown to facilitate the release of promoter-proximally paused Pol II (Rahl et al., 2010), enhance mRNA capping (Cowling and Cole, 2010), facilitate the transfers PAF1 to Pol II (Gerlach et al., 2017; Jaenicke et al., 2016), and enhance rate and processivity of transcription elongation by loading SPT5 onto Pol II (Baluapuri et al., 2019), and thereby support transcription of most active genes. In contrast, rapid depletion of MYC in leukemia and colon cancer cell lines affects transcription of a small subset of genes, suggesting that expression of a rather limited set of activated genes might depend on the presence of MYC (Muhar et al., 2018).

The activation of genes by MYC/MAX is antagonized by the MAX-Dimerization (MXD) proteins, MXD1-4, and MAX Network Transcriptional Repressor (MNT), henceforth collectively referred to as MXDs and MGA (Carroll et al., 2018). MXDs and MGA are bHLH transcription factors that form complexes with MAX and bind to the same E-boxes as MYC/MAX (Carroll et al., 2018; Conacci-Sorrell et al., 2014). MXDs recruit via their SID domain mSIN3-HDAC1/2 co-repressor complexes and repress transcription (Laherty et al., 1997; van Riggelen et al., 2010b).

Opposite to MYC, MXDs support cell cycle arrest and differentiation (Chen et al., 1995; Lahoz et al., 1994; Yang and Hurlin, 2017). Genetic studies in mice confirmed the antagonism between MYC and MXDs. *MXD1*^*-/-*^ mice show increased proliferation and detained differentiation of granulocyte precursors (Foley et al., 1998). Mice lacking *MXD2(MXI1)* display multiple histological abnormalities due to increased cell proliferation in several tissues, and are more susceptible to spontaneous and induced cancerogenesis (Schreiber-Agus et al., 1998). Depletion of MNT triggers increased cell proliferation (Hurlin et al., 2003; Nilsson et al., 2004). Mice bearing a deletion of *Mnt* in mammary glands develop spontaneous tumors with increased frequency, phenocopying transgenic overexpression of MYC (Toyo-oka et al., 2006). Finally, human *MNT, MXD1* and *MXD2* genes are located in regions that are frequently mutated in different cancer types (Cvekl et al., 2004; Edelmann et al., 2012; Schaub et al., 2018; Shapiro et al., 1994; Wechsler et al., 1994).

MYC, in particular in oncogenic or overexpressed conditions, has also the potential to repress transcription. The underlying mechanisms are not fully understood and a comprehensive set of bona-fide MYC-repressed genes is not known. This is in part due to the fact that MYC supports cell growth and proliferation, and thus, directly or indirectly promotes expression of all genes when compared to the transcription rates of resting cells. Hence, upon normalization lower than average activation of genes may appear as relative repression even though the genes are actually activated (Wolf et al., 2015). However, MYC has been shown to interact with the zinc-finger transcription factor MIZ1 (ZBTB17) and the related transcription factors, SP1 and YY1 (Poole and van Riggelen, 2017). MIZ1 regulates embryonic development and differentiation (Adhikary et al., 2003; Walz et al., 2014; Wolf et al., 2013; Wolf et al., 2015). MYC in association with MIZ1 has been shown to repress genes, including the cyclin-dependent kinase (CDK) inhibitor genes *p15* (*CDKN2B*), *p21* (*CDKN1A*) and *p27* (*CDKN1B*) and the circadian transcription factor genes *BMAL1* (*ARNTL*), *CLOCK* and *NPAS2* (Shostak et al., 2016; Staller et al., 2001; Walz et al., 2014; Wu et al., 2003; Yang et al., 2001). Mutations compromising the interaction of MYC with MIZ1 specifically affect the repressing but not the activating potential of MYC (Herold et al., 2002; Shostak et al., 2016; Si et al., 2010), indicating that MYC together with MIZ1 has the potential to directly repress transcription. In oncogenic conditions overexpressed MYC may recruit MIZ1 to a larger number of genes and attenuate their transcription (Lorenzin et al., 2016; Walz et al., 2014; Wolf et al., 2015). A low ratio of MYC versus MIZ1 occupancy (Lorenzin et al., 2016) and/or low relative affinity of promoters for MYC (de Pretis et al., 2017) appear to correlate with repression of transcription, suggesting that the ratio of activating MYC/MAX versus repressing MYC/MAX/MIZ1 complexes determines transcriptional outcome.

The development of lymphoma in mice is critically dependent on the interaction of MYC with MIZ1 (van Riggelen et al., 2010a). When the MYC/MIZ1 interaction is challenged by mutation, the repressive capacity of MYC is decreased, its pro-proliferative functions are reduced, and self-renewal of stem cells is compromised (Kerosuo and Bronner, 2016; Shostak et al., 2016).

In this study we addressed specifically the question whether and how MXDs have the potential to impact expression of genes that are repressed by MYC together with MIZ1. Using U2OS cells, which endogenously express MXDs, we report the surprising observation that MNT, MXD1, and MXD2 activate transcription of specific MYC-repressed genes, in parallel to their known function as transcriptional repressors of MYC-activated genes. We show that activation of transcription by MXDs relies on their physical interaction with MIZ1, and requires functional DNA binding and corepressor recruitment domains. We show that MXDs inhibit U2OS cell growth and proliferation through activation of MYC-repressed genes, and MXD-dependent activation of *p21* was particularly crucial.

## Results

### Endogenous MXDs activate the *p21* core promoter in U2OS cells

MXDs repress genes that are activated by MYC but a role of MXDs in the regulation of genes that are repressed by MYC has not been investigated. Since MXDs belong, like MYC, to the bHLH-ZIP family, we analyzed available ENCODE ChIP-seq data of MXD2 (GSM935498), MNT (GSE91968) and MYC (GSM822286, GSM822301). As expected, the analysis revealed that MXD2 and MNT binding overlaps with MYC binding (Fig. 1A). MXD2 and MNT were also recruited to MIZ1 binding sites in genes that are repressed by MYC such as *p21* and *VAMP4* (Fig. 1A). A genome-wide comparison of the cistromes of MNT and MYC (ENCODE data from MCF7 cells) and of MXD2 and MYC (ENCODE data from HELA cells) with the cistrome of MIZ1 in U2OS cells (Walz et al., 2014) revealed a significant overlap of MNT and MXD2 with MYC as well as with MIZ1 binding sites (Supplemental Fig. S1A). It should be noted that due to low sequence coverage we pooled binding sites of native MIZ1 from several ChIP-seq experiments published by Walz et al. Although the published ChIP-seq analyses are from different cell lines, the data suggest that MXDs and MYC might be recruited in a similar manner to genes that are co-occupied by MIZ1.

**Fig. 1.**
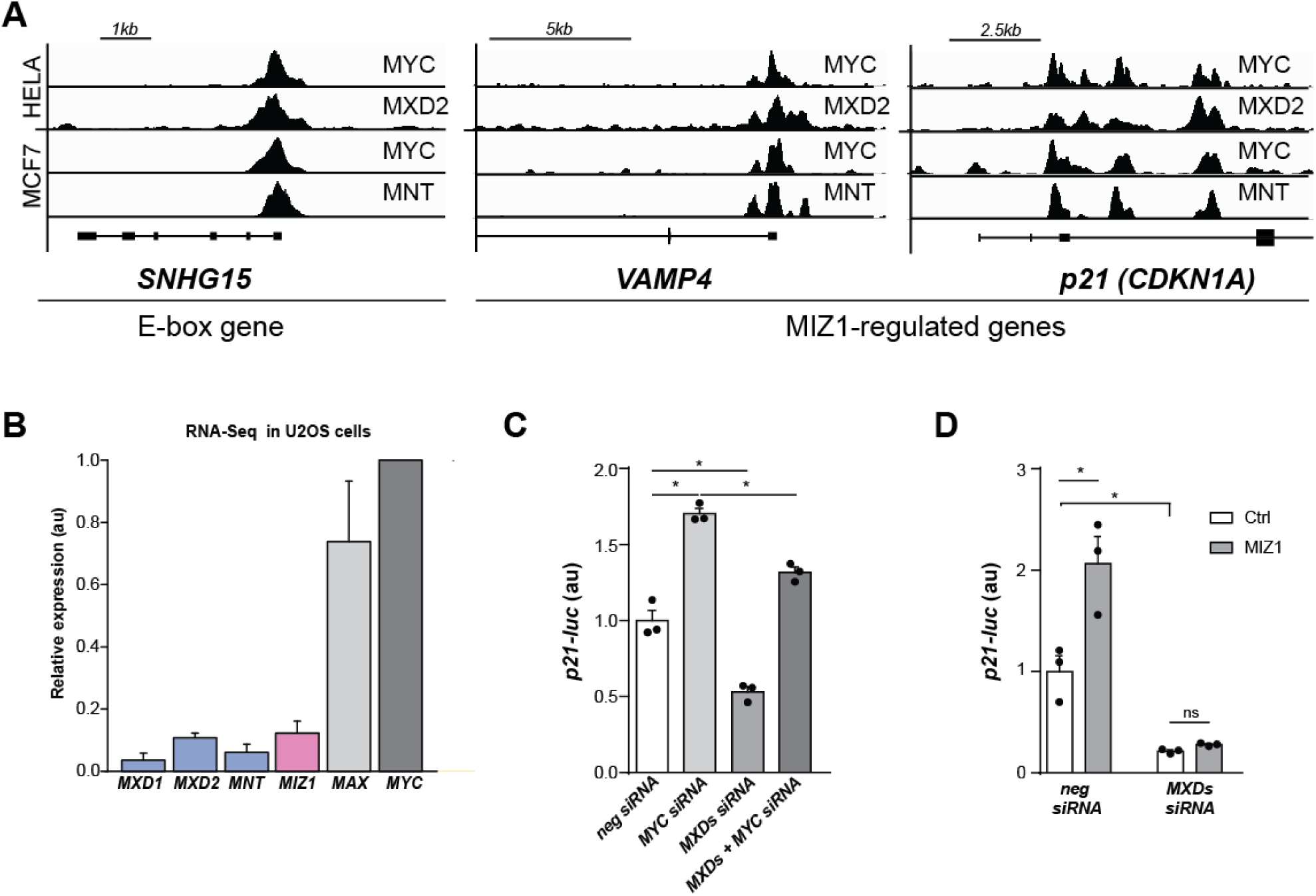
MXDs bind MIZ1/MYC sites and modulate MIZ1-dependent transcription. **(A)** *SNHG15, VAMP4*, and *p21 (CDKN1A)* loci with ChIP-seq signals of native MYC (GSM822286, GSM822301), MXD2 (GSM935498), and MNT (GSE91968) in HELA and MCF7 cells (based on data from ENCODE). **(B)** Relative expression (combined datasets GSM1632189, GSM1632191, GSM2341646, and GSM2341647 normalized to *MYC*) of *MXDs, MIZ1, MAX*, and *MYC* in U2OS cells quantified by RNA-seq (Elkon et al., 2015; Ibarra et al., 2016). It should be noted that MAX protein is about an order of magnitude more stable than MYC, and hence believed to be present in excess over its binding partners MYC and MXDs (Blackwood et al., 1992). **(C)** The ratio of MXDs versus MYC rather than their levels determine *p21-luc* expression. Quantification of bioluminescence (24 hours) from *p21-luc* reporter transfected in U2OStx cells pre-treated with siRNAs against *MXDs* (*MNT, MXD1*, and *MXD2*), *MYC* or *MIZ1*, as indicated. **(C)** MXDs support expression of *p21-luc*. Relative bioluminescence of *p21-luc* with and without induction of FLAG:MIZ1 (24 h) in U2OStx cells transfected with siRNAs against *MXDs* (*MNT, MXD1*, and *MXD2*) or negative siRNA (n=3). Data are presented as mean ± SEM. * P < 0.05; one-way ANOVA with Bonferroni post-test.

Expression of MXDs is tightly regulated and cell-type specific (Hooker and Hurlin, 2006). Available RNA-seq data indicate that MIZ1, MYC as well as its antagonists MXD1, MXD2, and MNT are expressed in U2OS cells (Fig. 1B) (Elkon et al., 2015; Ibarra et al., 2016). To assess whether MXDs are functional and impact expression of MYC repressed genes we analyzed expression of a *p21-luc* reporter. This luciferase reporter contains only the core promoter of the *p21* gene. Hence, it is highly likely that putative changes in *p21-luc* repression are directly due to regulation of transcription. *p21-luc* was previously shown to be repressed by MYC in MIZ1-dependent manner (Shostak et al., 2016; Wu et al., 2003). When U2OS cells were transfected with siRNAs against MYC the expression level of *p21-luc* was elevated (Fig. 1C), indicating that endogenous MYC had limited its expression. Surprisingly, expression of *p21-luc* was reduced when cells were treated with previously validated siRNAs against *MXDs* (Corn et al., 2005; Wu et al., 2012; Xu et al., 2007) (Fig. 1C and Supplemental Fig. S1B). The data demonstrate that MXDs are expressed at functional levels in U2OS cells, and that endogenous MXDs supported the expression of *p21-luc*. Expression of *p21-luc* in *MXD*-depleted cells was restored when cells were additionally transfected with siRNA against MYC (Fig. 1C). Together the data strongly suggest that the expression level of *p21-luc* in U2OS cells was established by the ratio of endogenous MYC versus MXDs.

We then produced U2OS cells overexpressing FLAG:MIZ1 in doxycycline-inducible fashion (Supplemental Fig. S1C) and measured expression of *p21-luc* (Fig. 1D). Expression of *p21-luc* increased when FLAG:MIZ1 was induced, indicating that MIZ1 was limiting in U2OS cells (Fig. 1D). The cells were then transfected with siRNAs against *MNT, MXD1* and *MXD2*. Downregulation of MXDs resulted in substantially reduced expression of *p21-luc* and overexpression of FLAG:MIZ1 failed to activate *p21-luc* when MXDs were depleted (Fig. 1D). The data indicate that both, endogenous and overexpressed MIZ1 required endogenous MXDs to activate *p21-luc* transcription.

### MXDs interact with MIZ1 to activate MYC-repressed genes

To analyze the impact of MXDs on MYC-repressed genes we generated stable U2OS cells expressing V5-tagged MXD1, MXD2, and MNT under control of a doxycycline-inducible promoter (see Supplemental Fig. S3B). Chromatin immunoprecipitation (ChIP) from these cells and from U2OS cells expressing MYC:V5 and FLAG:MIZ1, respectively, revealed that tagged MYC, MXDs and MIZ1 bound to the promoter of the *NCL* gene, which is a MYC activated gene, suggesting that the tagged TFs were functional in DNA binding (Fig. 2A). In agreement with the published ChIP-seq data shown in Fig. 1A, tagged MYC, MXDs and MIZ1 were also recruited the *p21* promoter (Fig. 2A), which is repressed by MYC.

**Fig. 2.**
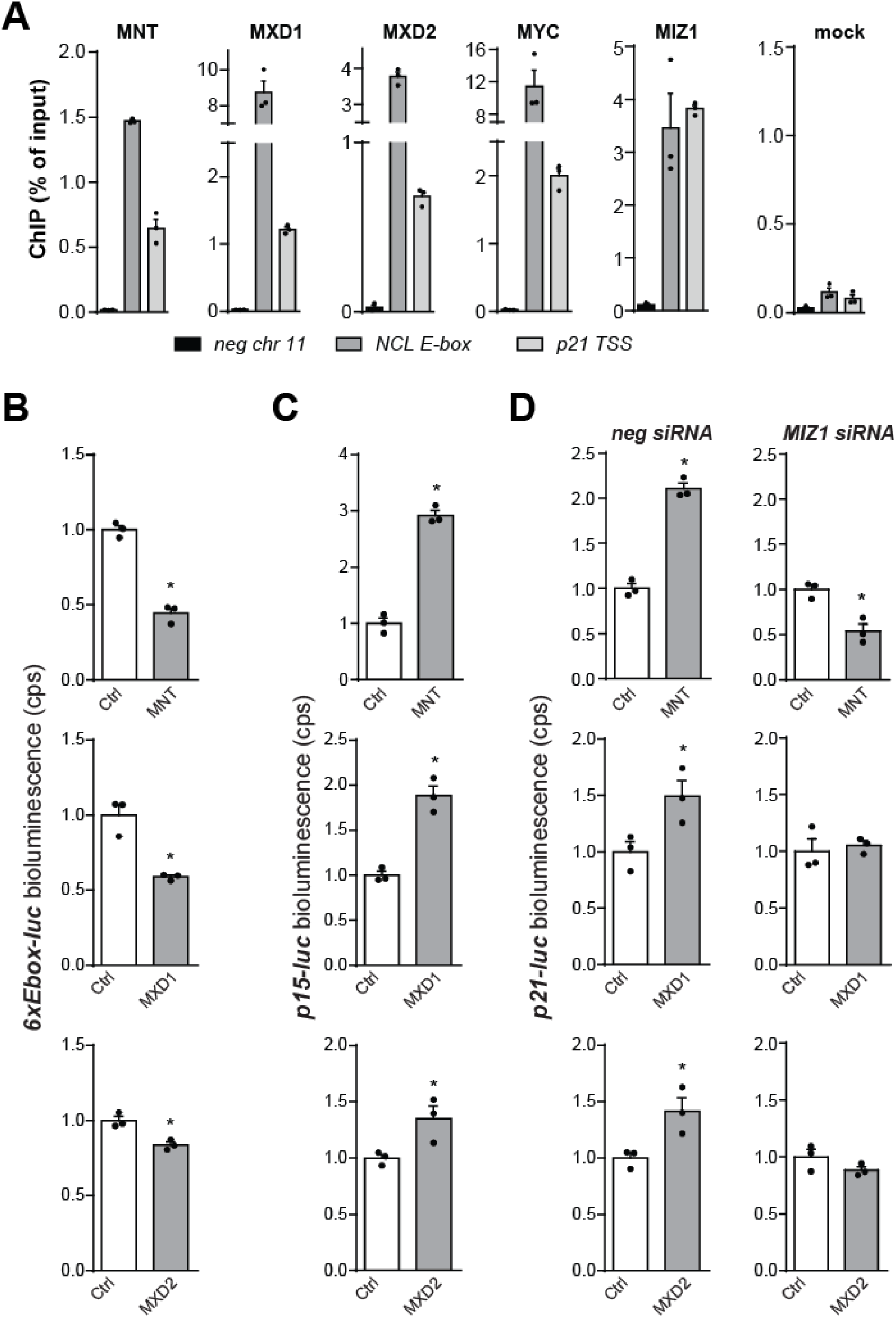
MNT, MXD1, and MXD2 repress E-box and activate MIZ1-target genes. **(A)** Binding (ChIP-PCR) of MNT, MXD1, MXD2, MYC, and MIZ1 to *NCL* and *p21*. Expression of the transcription factors in U2OStx cells was induced for 24 h with doxycycline (DOX) (n=3). Bioluminescence of *6xEbox-luc* **(B)** and *p15-luc* **(C)** reporters from U2OStx tetO-MNT, MXD1, and MXD2 cells after induction with doxycycline (n=3). Cells were transiently transfected with luciferase plasmids and bioluminescence from the living culture was measured 24 h after DOX induction. **(D)** Activation of *p21-luc* in U2OS cells overexpressing MNT, MXD1, and MXD2 after induction with DOX (24 h). Cells pre-treated with siRNA were transfected with reporter plasmid one day before DOX induction (n=3). Data are presented as mean ± SEM. * P < 0.05; Student’s *t*-test.

We have previously shown that doxycycline-induced overexpression of MYC supported expression of a *6xEbox-luc* reporter and inhibited expression of *p15-luc* and *p21-luc* reporters (Shostak et al., 2016). Doxycycline-induced overexpression of MNT, MXD1, and MXD2 attenuated expression of a *6xEbox-luc* while they supported elevated expression of *p15-luc* and *p21-luc* reporters (Fig. 2B, C, and D). Activation and repression of the reporters were strongest for MNT and rather moderate for MXD2, yet even these moderate increases in expression were significant and highly reproducible. Interestingly, MNT, MXD1, and MXD2 did not support elevated expression of *p21-luc* when MIZ1 was depleted by siRNA (Fig. 2D and Supplemental Fig. S2A). These data indicate that activation of the reporter required endogenous MIZ1.

In accordance with the reporter assays, MXDs also impacted the expression levels of endogenous genes. Thus, induction of MNT, MXD1, and MXD2, attenuated expression of *NCL* and *SNHG15*, confirming the repressive role of MXDs on MYC-activated genes.

In contrast, MNT, MXD1, and MXD2 supported expression of endogenous *p15, p21, p27*, and *CEBPA* (Supplemental Fig. S2B). As observed with the gene reporters, MNT was the strongest activator under our conditions while the activating potential of MXD2 was rather modest. The activation of *p15* by MXDs was pronounced while the activation of *p27* was significant but rather small. Together the reporter assays and the measurement of the expression levels of the corresponding endogenous genes indicate that overexpressed MXDs have the potential to activate in U2OS cells the cyclin-dependent inhibitor genes *p15* and *p21*, and *p27* as well as *CEBPA*. The extend of activation by the individual MXDs differs between genes and is dependent on MIZ1. Furthermore, overexpression of ectopic MXDs and downregulation of endogenous MXDs, respectively, have the opposite effect on *p21-luc* expression. Thus, our data indicate that MXDs (endogenous and ectopic) have the potential to activate these MYC-repressed genes in U2OS cells under conditions where they are active as repressors of MYC-activated genes.

The above measurements were done with confluent U2OS cells, which are growing rather slowly. We chose these conditions to avoid or minimize potential indirect effects on gene expression that could be associated with MXD-dependent differences in cell number (due to growth or apoptosis). Colorimetric cell counting by WST-8 staining indicated that induction of MXDs had little impact on cell growth and viability under such conditions (Supplemental Fig. S2C), and quantification of *GAPDH* expression from the entire cultures confirmed the results of the WST-8 assay (Supplemental Fig. S2D). Hence, the MXD-dependent increase in expression levels of *p15, p21*, and *p27* (*luc*-reporters and endogenous genes) is due to transcriptional regulation.

### MXDs interact with MIZ1 and directly activate MYC-repressed genes

The bHLH domains of MYCs and MXDs are highly conserved (Fig. 3A) (Hurlin et al., 1997). In order to analyze whether MXDs physically interact with MIZ1, we expressed in HEK293 cells V5 tagged versions of MXD1, MXD2, and MNT together with FLAG-tagged MIZ1. Pull-down assays revealed that FLAG-tagged MIZ1 formed complexes with MXD1, MXD2, and MNT (Fig. 3B and Supplemental Fig. S3A). We then set out to generate MXD mutants that are compromised in their ability to interact with MIZ1. The interaction of MYC with MIZ1 is critically dependent on V393 and V394 in the bHLH domain of MYC (Herold et al., 2002; Shostak et al., 2016). A sequence comparison revealed that V393 is conserved in MYC proteins, while MXDs carry a leucyl residue in the corresponding position (Fig. 3A). To assess whether MXDs interact with MIZ1 via a similar interface as MYC we expressed in HEK293 cells V5-tagged versions of MNT, MXD1, and MXD2 with leucyl to aspartyl (L-D) substitutions in the position corresponding to V393 of MYC. Subsequent pull-down assays revealed that the mutation abolished the interaction of MNT_L258D_ with MIZ1 (Fig. 3C). MXD1_L95D_, and MXD2_L106D_ were substantially compromised in their capacity to interact with MIZ1 (Fig. 3D and Supplemental Fig. S3B). The potential of the mutant MXDs to bind MAX was not affected (Supplemental Fig. S3C). The data suggest that MYC and MXDs interact with MIZ1 in corresponding manner.

**Fig. 3.**
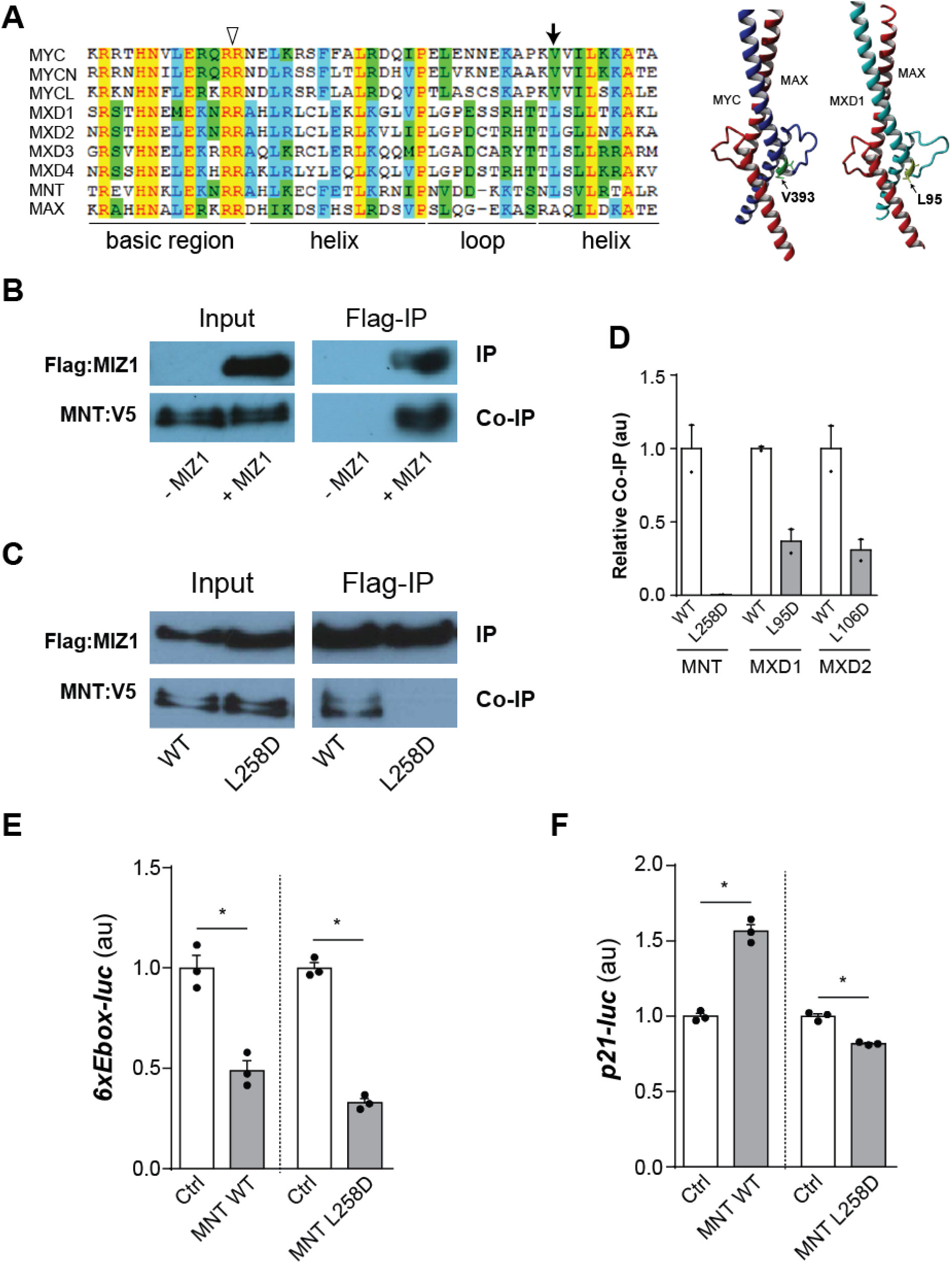
MNT, MXD1, and MXD2 bind MIZ1 via a conserved interface. **(A)** Left: Alignment of bHLH domains of MYC (MYC, MYCN, MYCL) and MXD (MXD1, MXD2, MXD3, MXD4, and MNT). The position corresponding to V393 of MYC is indicated by an arrow. The open arrowhead indicates position of two conserved arginyl residues (RR). Right: Structure of MYC/MAX and MXD1/MAX dimers with indicated positions of V393 and L95, respectively (Nair and Burley, 2003). **(B)** Co-immunoprecipitation of V5-tagged MNT with FLAG-tagged MIZ1 from HEK293 lysates. **(C)** FLAG:MIZ1 pulldown and co-immunoprecipitation of WT and L258D versions of MNT expressed in HEK293 cells. MIZ1 and MNT were tagged with FLAG and V5 epitopes, respectively. **(D)** The efficiency of co-immunoprecipitation of WT and L-D versions of MXDs was quantified by densitometry (n=2). Expression of *6xEbox-luc* **(E)** and *p21-luc* **(F)** reporters in U2OStx cells after DOX induction (24 h) of WT or L-D versions of MXDs (n=3). Ctrl: non-induced control. Data are presented as mean ± SEM. * P < 0.05; one-way ANOVA with Bonferroni post-test.

We then generated U2OS cells expressing doxycycline inducible L-D versions of MXDs (Supplemental Fig. S3D). The capacity of MNT_L258D_, MXD1_L95D_, and MXD2_L106D_ to repress *6xEbox-luc* was not affected (Fig. 3E and Supplemental Fig. S3E), indicating that the mutant proteins were functional repressors. However, the L-D versions of MXDs did not support elevated expression *p21-luc* (Fig. 3F and Supplemental Fig. S3F), suggesting that the capacity to activate this promoter was dependent of their interaction with MIZ1. Furthermore, MNT_L258D_, MXD1_L95D_, and MXD2_L106D_ attenuated expression of endogenous *NCL* and *SNHG15* genes as efficiently as the corresponding WT proteins but showed reduced activation of *p15, p21, p27*, and *CEBPA* genes (Supplemental Fig. S3G). Together the data suggest that activation of these MYC-repressed genes by MXDs was supported by their interaction with MIZ1. The data are not compatible with an indirect activation of transcription by sequestration of MAX and thereby relieving MYC-dependent repression.

### MYC and MXDs require DNA binding to modulate MIZ1 activity

To test whether MYC and MXDs require an intact bHLH domain in order to associate with MIZ1-target genes, we replaced in the basic region of the DNA binding domain two critical neighboring arginyl residues (RR) by aspartyls (DD) (see Fig. 3A). Recruitment of the corresponding mutants, MYC_RR367DD_, MNT_RR232DD_, MXD1_RR68DD_, and MXD2_RR79DD_, to MYC-activated and MYC-repressed genes was reduced (Fig 4A). However, the RR-to-DD substitutions in the DNA-binding domains of MYC and MXDs did not affect their ability to interact with MIZ1 in a pull-down assay (Supplemental Fig. S4A).

**Fig. 4.**
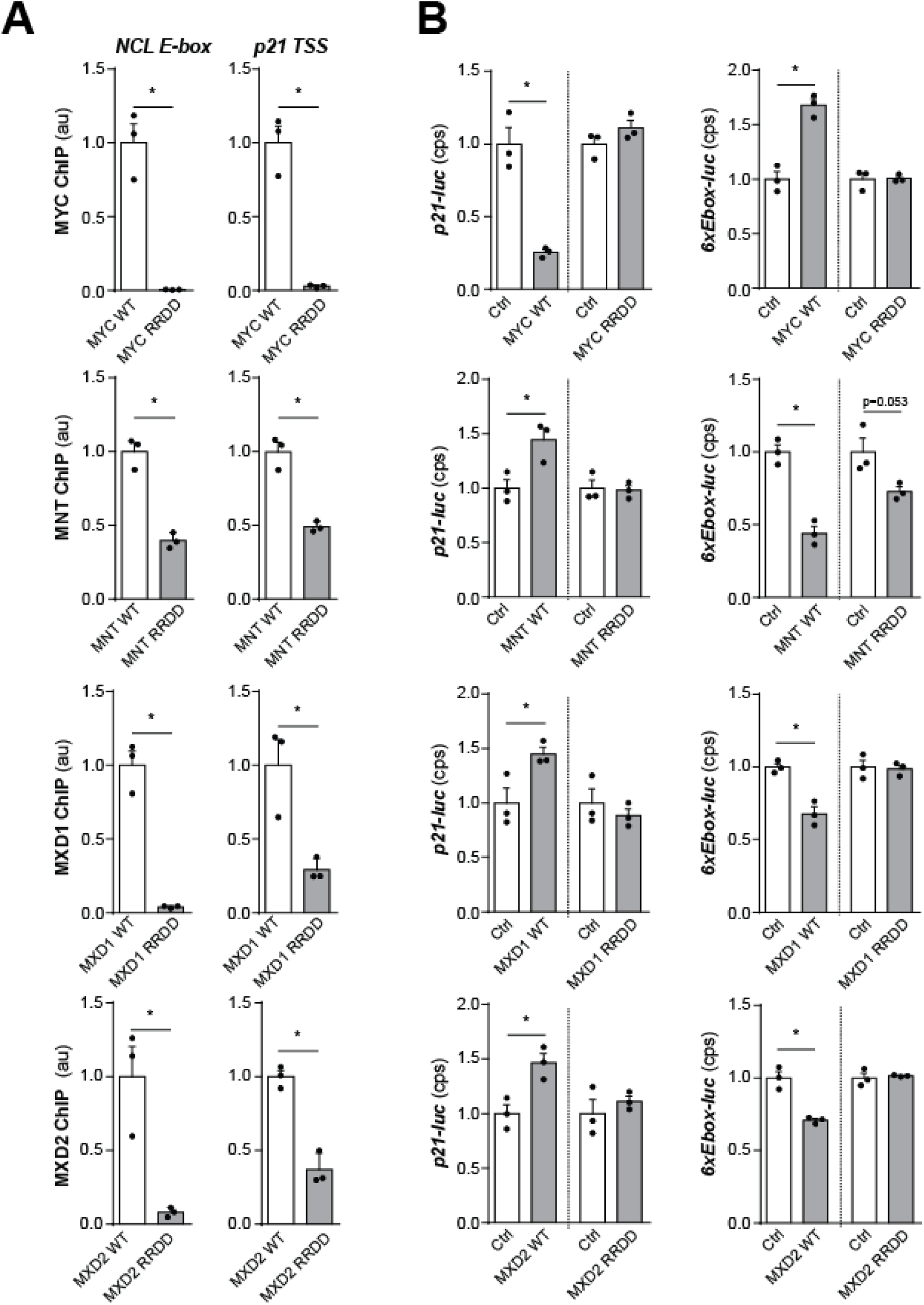
MYC and MXDs require DNA binding to regulate MIZ1 targets. **(A)** Binding of MYC and MXDs to DNA requires an intact bHLH domain. ChIP-PCR of MYC, MNT, MXD1, and MXD2 (WT and RRDD versions) at promoters of *NCL* and *p21* in U2OStx cells 24 h after DOX induction (n=3). * P < 0.05; Student’s t-test. **(B)** Transcription regulation by MYC and MXDs requires an intact bHLH domain. Bioluminescence of *p21-luc* and *6xEbox-luc* reporters measured from U2OStx cells 24 hours after DOX induction of WT and RRDD versions of MYC and MXDs (n=3). Ctrl: non-induced control. * P < 0.05; one-way ANOVA with Bonferroni post-test. Data are presented as mean ± SEM.

Overexpression of MYC_RR367DD_ in U2OS cells (Supplemental Fig. S4B) neither activated *6xEbox-luc* nor repressed *p21-luc* (Fig. 4B) (Shostak et al., 2016), suggesting that the DNA-binding domain of MYC is required for both, activation and repression of genes. Similarly, RR-to-DD substitutions in the DNA binding domains of MXDs impaired their potential to repress *6xEbox-luc* and their capacity to activate *p21-luc* (Fig. 4B). These data suggest that the DNA-binding domains of MXDs are required for repression and activation of genes. Since the RR-to-DD substitutions did not affect the interaction of MXDs with MIZ1, the data also indicate that MXDs did not indirectly activate *p21-luc* by squelching MIZ1 away from MYC/MIZ-repressed promoters.

MXDs harbor a SID domain by which they recruit co-repressor complexes (Laherty et al., 1997). The SID domains of MXDs are short N-terminal segments of about 20 amino acid residues (aa). We deleted the SID domains of MNT (Δaa 2-16), MXD1 (Δaa 2-20), and MXD2 (Δaa 2-20) (Supplemental Fig. S5). It has been previously shown that deletion of the SID domain does not affect the interaction of MXDs with MAX (Ayer et al., 1995). ChIP analysis revealed that binding of MXDΔSID mutants to the promoters of *NCL* and *p21* was not compromised (Fig. 5A). Previous data (Hurlin et al., 1997) had shown that MXDΔSID mutants failed to repress MYC-activated genes. In agreement with these data MXDΔSID mutants failed to repress the *6Ebox-luc* reporter in HEK293 cells (Fig. 5B). MNTΔSID even activated *6Ebox-luc.* Surprisingly, however, the ΔSID versions of MXDs were compromised in their ability to activate *p21-luc* (Fig. 5C), suggesting that the SID domain is also required for MIZ1-dependent transactivation of MYC-repressed genes.

**Fig. 5.**
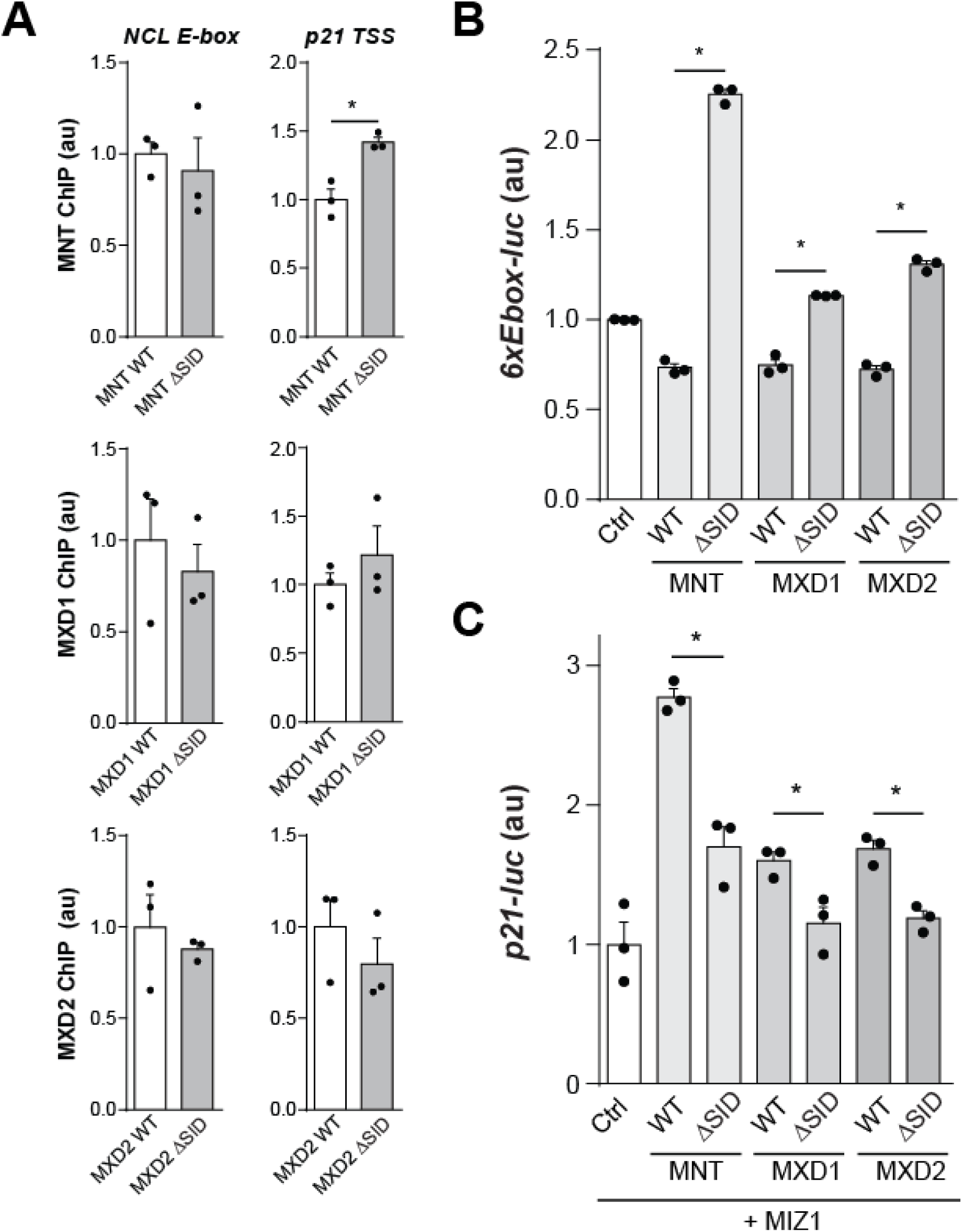
MXDs require SID domains to activate and repress genes. **(A)** DNA binding of MXDs does not require SID domains. ChIP-qPCR analysis of showing that DOX-induced (24 hours) WT and ΔSID versions of MNT, MXD1, and MXD2 bind to *NCL* and *p21* genes in U2OStx cells (n=3). **(B**,**C)** The SID domains of MXDs are required for repression of *6xEbox-luc* and activation of *p21-luc.* **(B)** Luciferase reporter assay of *6xEbox-luc* in HEK293 cells transfected with wild type or ΔSID MNT, MXD1, and MXD2 (n=3). **(C)** *p21-luc* expression in HEK293 cells transfected with MIZ1 and co-transfected with wild type or ΔSID versions of MXDs (n=3). * P < 0.05; one-way ANOVA with Bonferroni post-test. Data are presented as mean ± SEM.

### MXDs inhibit cell growth in MIZ1-dependent manner

Expression of MXDs reduces cell growth and proliferation in various cellular models (Chin et al., 1995; Delpuech et al., 2007; Hurlin et al., 1997). Doxycycline-induced overexpression of MXDs also reduced proliferation of growing U2OS cells that were seeded at low density (Fig. 6A). To analyze whether MXDs inhibit growth in MIZ1-dependent fashion we overexpressed the MIZ1-interaction mutants, MNT_L258D_, MXD1_L95D_, and MXD2_L106D_, which are functional repressors of MYC-activated genes (see Fig. 3E and Supplemental Fig. S3E, G). The capacity of MNT_L258D_, MXD1_L95D_, and MXD2_L106D_ to inhibit cell growth and proliferation was severely blunted (Fig. 6A and Supplemental Fig. S6A), suggesting that inhibition of U2OS cell growth and proliferation by MXDs relies critically on their interaction with MIZ1.

**Fig. 6.**
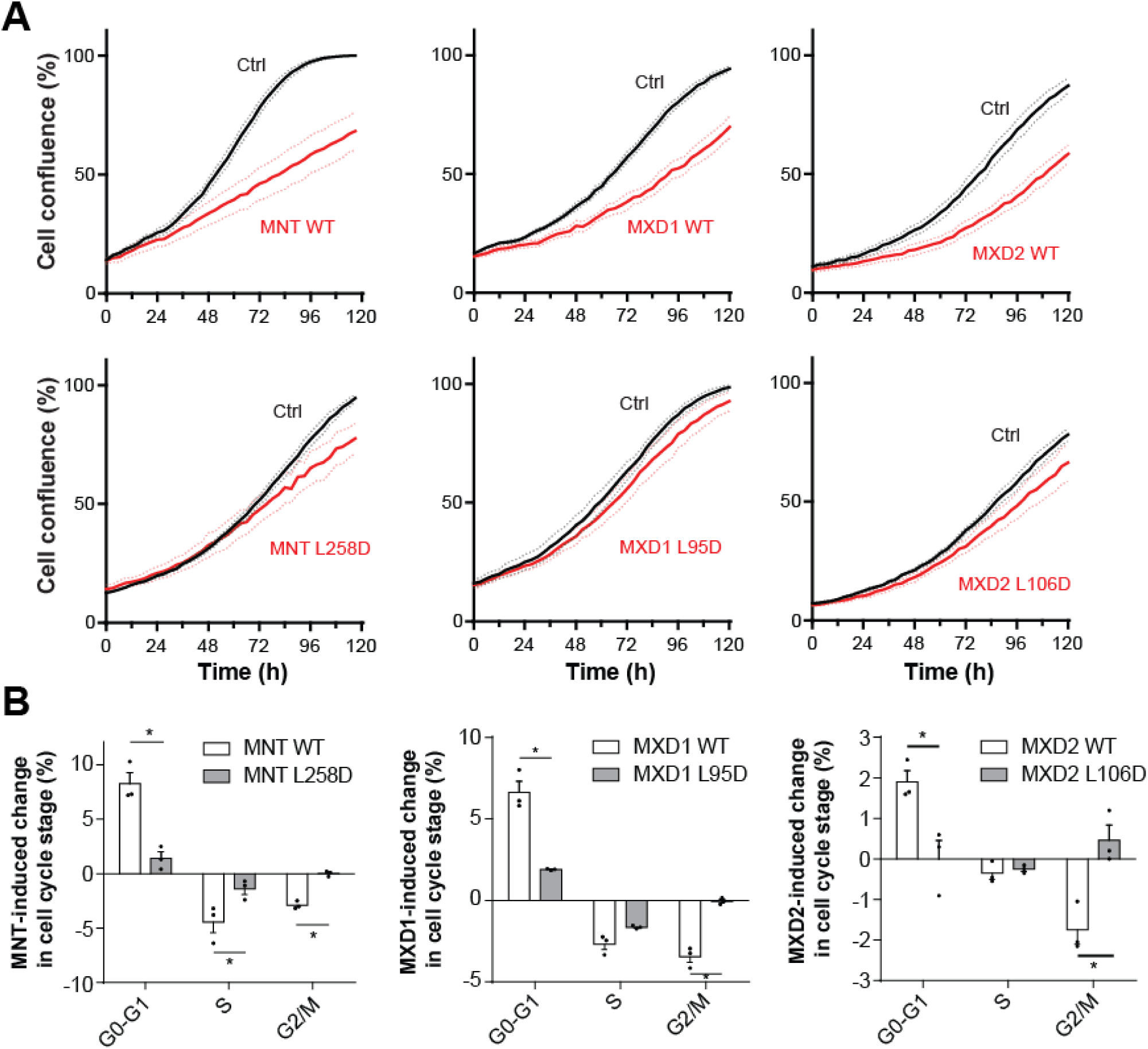
MXDs attenuate cell growth in MIZ1-dependent manner. **(A)** L-D versions of MXDs do not inhibit proliferation. Proliferation of U2OStx cells expressing DOX-induced WT and L-D versions MXDs. Confluence of cells, treated with DOX (red curves) or PBS (Ctrl, black curves), was recorded with an Incucyte ZOOM system (n=3). **(B)** Wild type but not L-D mutants of MXDs attenuate cell proliferation. Cells were treated with DOX or PBS for 48 h and then were stained with propidium iodide and cell cycle stage was analyzed by FACS. Changes of the fractions of cells in G0-G1, S, and G2 triggered by expression of WT or L-D alleles of MXDs are shown (n=3). * P < 0.05; two-way ANOVA with Bonferroni post-test. Data are presented as mean ± SEM.

We then asked whether the reduced apparent kinetics of cell growth was due to an increase in apoptosis of U2OS cells or a reduction in cell cycle frequency. Cell cycle stage-specific FACS-sorting of propidium iodide stained cells revealed that overexpression of MXDs reduced the fraction of cells in G2 and M phases, while overexpression of L-D mutants of MXDs had no significant effect on the cell cycle (Fig. 6B). Measurement of caspase activity revealed that cells gradually started to undergo apoptosis only after about 60 h of growth, long after cells reached confluence (Supplemental Fig. S6B). Overexpression of MXDs reduced apoptosis while overexpression of MYC increased apoptosis, as reported previously (Herkert et al., 2010) (Supplemental Fig. S6B). Together the data demonstrate that MXDs inhibit proliferation of U2OS cells by inhibiting the cell-cycle.

Finally, we addressed the role MXDs in regulating growth of U2OS cells via *p21*. When *p21* was depleted in growing U2OS cells the growth rate did not further increase, suggesting that growth was limited by other factors. Overexpression of MXDs attenuated proliferation of U2OS cells but failed to do so when MXD-induced accumulation of *p21* was prevented by siRNA treatment (Supplemental Fig. S7C, D). Hence, the induction and/or repression of other genes by MXDs could obviously not compensate for the lack of *p21*. These data indicate that regulation of the CDK inhibitor gene, *p21*, is a major pathway by which MXDs and MIZ1 impact proliferation of U2OS cells.

## Discussion

MXD proteins encompass a group of transcriptional repressors that antagonize the activation of genes by MYC (Conacci-Sorrell et al., 2014). The aim of this work was to investigate whether and how MXDs impact on such genes that are not activated but repressed by MYC. We therefore focused specifically on the regulation of a selected group of gene promoters (mainly *p15, p21*, and *p27*) that were previously shown by various means to be directly repressed by MYC in MIZ1-dependent manner (Shostak et al., 2016; Si et al., 2010; Staller et al., 2001; Walz et al., 2014; Wu et al., 2003; Yang et al., 2001), and we restricted our analyses on U2OS cells, which express MXDs as well as MYC at functional levels. We report the unexpected and surprising finding that MXDs activated transcription of this validated group of MYC-repressed genes. Thus, the antagonizing role of MXDs appears to extends to the limb of genes that are repressed by MYC, i.e., gene that are activated MYC are repressed by MXDs, and vice versa, genes that are repressed MYC are activated by MXDs. Since the activating function of MYC is antagonized by MXDs is seems conceivable that also the repressing function of MYC requires regulatory counterbalance, in particular since MYC-repressed genes include crucial inhibitors of the cell cycle.

Several lines of evidence support that transcription of genes such as *p15, p21*, and *p27* was directly activated by MXDs. Most importantly, MXDs are functionally expressed in U2OS cells and MXD downregulation resulted in elevated expression of the selected MYC-repressed genes. We controlled WST-8 staining that the apparent induction of these MYC-repressed genes was not artificially due to differences in number and size of living cells (biomass) in MXD-induced versus control-treated cultures (e.g. by less apoptosis of MXD-induced cells).

In addition to the analysis of endogenous gene expression we measured the impact of MXDs on luciferase reporter genes to assess the transcriptional regulation of the core promoters. The *p21-luc* reporter contains a short, truncated core promoter with binding sites for SP1, MIZ1, and MYC (Wu et al., 2003), and also for MXDs. Thus, analysis of *p21-luc* minimizes the potential impact of other transcription factors implicated in the complex regulation of the endogenous *p21* gene. Induction of MXDs in confluent cultures of U2OS cell triggered an increase in *p21-luc* bioluminescence measured from the entire culture while the number of viable cell did not increase relative to control cultures. Moreover, depletion of endogenous MXDs had the opposite effect (reduced expression of *p21-luc*) than overexpression of MXDs (elevated expression of *p21-luc*). Together, our data show that MXDs have the potential to activate in U2OS cells transcription of selected MYC-repressed genes.

We then analyzed the molecular basis underlying the MXD-dependent transcription activation. These analyses were based on functional comparison of overexpressed WT MXDs with mutant MXD versions defective in MIZ1 interaction, DNA binding and co-repressor recruitment, respectively. In principle, overexpressed MXDs could induce MYC-repressed genes indirectly by sequestering MAX, and thereby reducing the levels of repressive MYC complexes. However, DNA-binding mutants as well as MIZ-interaction mutants of MXDs interacted with MAX as efficiently as WT MXDs but did not induce MYC-repressed genes, which is not compatible with indirect gene activation by sequestration of MAX.

Potentially MXDs could (via the canonical pathway) repress putative MYC-activated genes that encode (co)repressors of MYC-repressed genes, and thereby indirectly, i.e. by repression of a (co)repressor, induce transcription. However, the MIZ interaction mutants of MXDs were active as repressors of MYC-activated genes but failed supporting expression of the selected MYC-repressed genes, excluding indirect activation via repression of a (co)repressor.

Together, our data suggest that MXDs activated MYC-repressed genes in direct manner in addition to their function as repressors of MYC-activated genes. The dual function of MXDs is not unprecedented as MYC itself also acts as activator and repressor, and interacts via its MYC-boxes with factors associated with transcription activation and repression (Baluapuri et al., 2019; Kress et al., 2015; Poole and van Riggelen, 2017; Tu et al., 2018). The conditions and mechanisms specifying MYC as either (general) activator or repressor of particular genes are only partly understood.

MXDs repress MYC-activated genes by recruiting via their short (∼20 aa) N-terminal SID domains mSIN3-HDAC co-repressor complexes (Laherty et al., 1997) but MXDs do not contain known domains for the recruitment of co-activators. SID deletions do not affect the recruitment of MXDs to their target genes but compromised repression of *6xEbox-luc*, and, surprisingly, also activation of *p21-luc*. The underlying mechanism remains obscure and awaits further investigation.

Together, these data indicate that MXDs (at physiological and overexpressed levels in U2OS cells) have the potential to directly activate in MIZ1-dependent fashion selected genes that are repressed by MYC/MIZ1. While MIZ1 may activate transcription by default, it is limiting in U2OS cells and its regulatory potential in the presence of endogenous levels of MXDs and MYC is determined by the functional ratio of MXDs versus MYC. In growing U2OS cells MYC may dominate and hence MIZ1 is predominantly repressing, while in non-growing U2OS cells MDXs may functionally dominate and MIZ1 is predominantly activating. The physiological relevance and contribution of MXDs to the expression of MYC-repressed in the context of a living organism is beyond the scope of this manuscript and remains to be investigated.

MYC and MXDs are regulators of cell proliferation. We show here that overexpression of MXDs attenuated proliferation of growing U2OS cells (non-confluent cultures), consistent with their reported role as antagonists of MYC, which stimulated proliferation under corresponding conditions. Mutant versions of MXDs with reduced affinity for MIZ1 were compromised in their capacity to inhibit cell proliferation, just as a corresponding MIZ1-interaction mutant of MYC was recently shown to be compromised in its capacity to stimulate cell proliferation (Kerosuo and Bronner, 2016; Shostak et al., 2016; van Riggelen et al., 2010a). Hence, at least in U2OS cells MXDs antagonized MYC’s pro-proliferative function predominantly in MIZ1-dependent manner via the regulation of MYC-repressed genes. Overexpression of MXDs failed to attenuate cell growth when expression of *p21* was suppressed by siRNA, indicating that regulation of *p21* was of particular importance in U2OS cells.

In summary, our results reveal that transcription factors of the MXD family, which are characterized repressors of MYC-activated genes, show transcription activating properties at genes that are repressed by MYC together with MIZ1. Thus, our findings suggest that activation and repression of genes by MYC could be coordinated by a reciprocal antagonism of MXD proteins.

## Materials and Methods

### Cell culture and transfections

U2OStx and HEK293 cells (Shostak et al., 2016) were maintained in DMEM with 10% FBS and 1x PenStrep. Cell culture reagents were obtained from Life Technologies unless indicated differently. Inducible U2OStx cells overexpressing different transgenes were obtained by stable transfection with *AhdI*-linearized pcDNA4/TO vector using Xfect (Clontech). Resistant clones were selected with 50 µg/ml hygromycin and 100 µg/ml zeocin (Invivogen) for 2 weeks and pooled together. For siRNA transfections, U2OS cells were seeded on 24 or 96 well plates and next day transfected with the siRNAs (sequences are given in Supplemental Table 1) using Lipofectamine RNAiMAX reagent. Cells were kept in the siRNA transfection mix minimum 24 hours before further applications. For luciferase reporter assays, HEK293 cells were transfected with the indicated plasmids using Lipofectamine2000. Next day, luciferase expression was measured using Dual-luciferase Reporter Assay (Promega) and an EnSpire Reader (Perkin Elmer).

### Plasmid constructs

*6xEbox-luc, p15-luc*, and *p21-luc* reporters were used previously (Shostak et al., 2016). Vectors containing the ORFs of *MNT* and *MXD1* were kindly provided by Prof. Bernhard Lüscher. The *MXD2* ORF was amplified from U2OS cDNA. To produce inducible constructs V5-tagged ORFs of *MXDs* were cloned in pcDNA4/TO vector. Subsequent L-D, RRDD, and ΔSID mutagenesis, respectively, was performed using DF-Pfu polymerase (Bioron). Cloning and mutagenesis primers are available upon request.

### Bioluminescence measurements

For bioluminescence measurements, transgenic U2OStx cells, untreated or pretreated with siRNA, were transiently transfected with luciferase reporters using Xfect (Clontech). After 24 hours the growth medium was replaced with prewarmed luminescence medium (DMEM w/o Phenolred (Cat. no. 21063-029) supplemented with 10% FBS, 1x Normocin (Invivogen), and 0.125 µM luciferin (BioSynth), 10 ng/ml doxycycline or PBS) and plates were measured with an EnSpire Reader (Perkin Elmer).

### Gene expression analysis

Total RNA from U2OStx cells was extracted with TriFaster (GeneON) and cDNA synthesis was performed with Maxima First Strand cDNA Synthesis Kit (Thermo). qPCR was performed on LightCycler 480 (Roche) using Maxima SYBR Green/ROX qPCR Master Mix (Thermo) and gene expression was quantified using a ΔΔCt method relative to *GAPDH*. Primer sequences are listed in Supplemental Table 1.

### ChIP

U2OStx cells overexpressing different transgenes were incubated with doxycycline for 24 h and then cross-linked in 1% formaldehyde for 10 min. Chromatin was prepared as described previously (Shostak et al., 2016). Sheared chromatin was incubated overnight at 4°C with 40 µl of salmon sperm DNA-blocked anti-FLAG (A2220, Sigma-Aldrich) and anti-V5 (A7345, Sigma-Aldrich) beads. Subsequently, precipitated chromatin was washed and recovered as previously described (Shostak et al., 2016). Samples were then analyzed by qPCR, and values were normalized to percentage of input. Primer sequences are listed in Supplemental Table 1.

### Cell confluence and fluorescent microscopy

For proliferation assays, siRNA-transfected or untreated transgenic U2OStx cells were seeded on transparent 96 well plates (∼3000 cells per well) and next day induced with standard growth medium containing 10 ng/ml doxycycline or PBS for control. To quantify apoptosis, growth medium was supplemented with Caspase-3/7 Green Reagent (4440, Essen Bioscience). Cell confluency and apoptosis were measured with an IncuCyte ZOOM reader (Essen Bioscience) using in-built software.

### Co-immunoprecipitation and Western blotting

Protein lysates of U2OS cells were prepared by incubation with ice cold lysis buffer (Shostak et al., 2016) and subsequent sonication in the ultrasonic bath (Merck) for 10 min. Pre-cleared lysates (centrifugation at 16000xg for 10 min at 4°C) were boiled with 4x Laemmli buffer, separated using 12% SDS-PAGE and transferred on nitrocellulose membranes. Membranes were decorated with anti-FLAG (1:5000, M2, Sigma-Aldrich), anti-V5 (1:5000, 46-0705, Life Technologies), and anti-Tubulin (1:1000, WA3) antibodies in TBS 5% milk at 4°C overnight. Next day, membranes were incubated with respective HRP-conjugated secondary antibodies and exposed to X-ray films. For co-IP experiments, transfected HEK293 cells were collected and prepared as described above. Then cell lysates (500 µg total protein) were incubated with 40 µl of PBS-washed anti-FLAG M2 beads (Sigma-Aldrich) at 4°C overnight. Next day beads were washed 3 times with PBS supplemented with 500 mM NaCl and 1% Triton X100 and precipitated proteins were eluted by boiling in 4x Laemmli buffer and loaded on 12% SDS-PAGE gels.

## Acknowledgments

We thank Bernhard Lüscher (RWTH Aachen University) for plasmid constructs and Bianca Ruppert and Maria Luisa Möller-Winheim for technical assistance. The work was supported by the Collaborative Research Centre TRR186 of the Deutsche Forschungsgemeinschaft (DFG). MB is a member of CellNetworks.

## Author contributions

AS performed the experiments and GS did the bioinformatics analysis. AS, AD and MB planned and designed the experiments and wrote the manuscript.

