## Supplementary Figures for "MXD/MIZ1 complexes activate transcription of MYC-repressed genes"

Fig. S1, Shostak et al

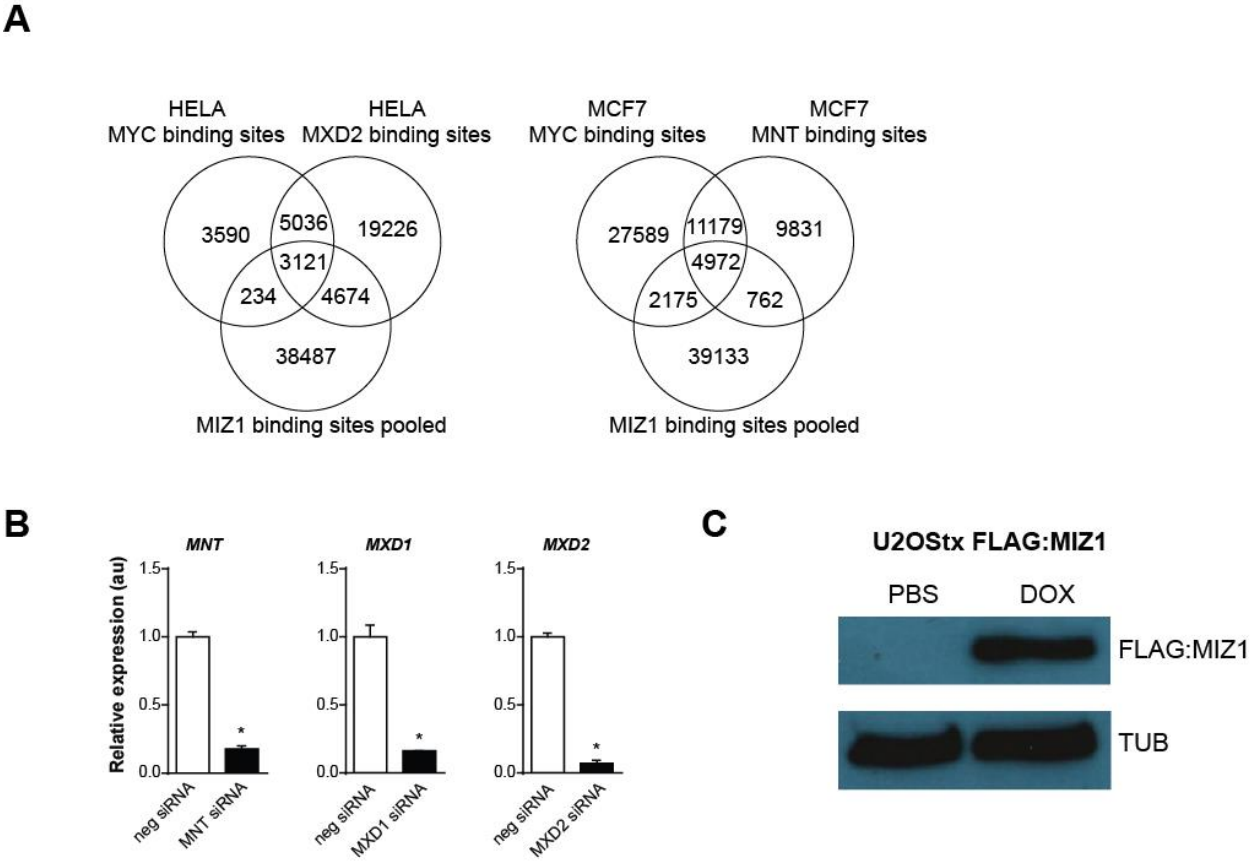


**Supplemental Figure S1. (A)** Venn diagram showing the overlap between native MYC (GSM822286 for HELA and GSM822301 for MCF7), MIZ1, and MNT (GSE91968) or MXD2 (GSM935498) binding sites in MCF7 and HELA cells, respectively. MIZ1 binding sites (GSM1088664, GSM1231600, GSM1231599) were pooled from several ChIP-seq experiments (Walz et al. 2014). **(B)** Downregulation of MXDs by siRNA. qPCR quantification of MNT, MXD1, and MXD2 transcripts in U2OStx cells transfected with the respective siRNAs (n=3). Total RNA was isolated 24 h after siRNA transfection. Data are presented as mean ± SEM. * $P < 0.05$; Student's t-test. **(C)** Western blot analysis of U2OStx cell lysates expressing DOX-induced FLAG:MIZ1.

Fig. S2 Shostak et al

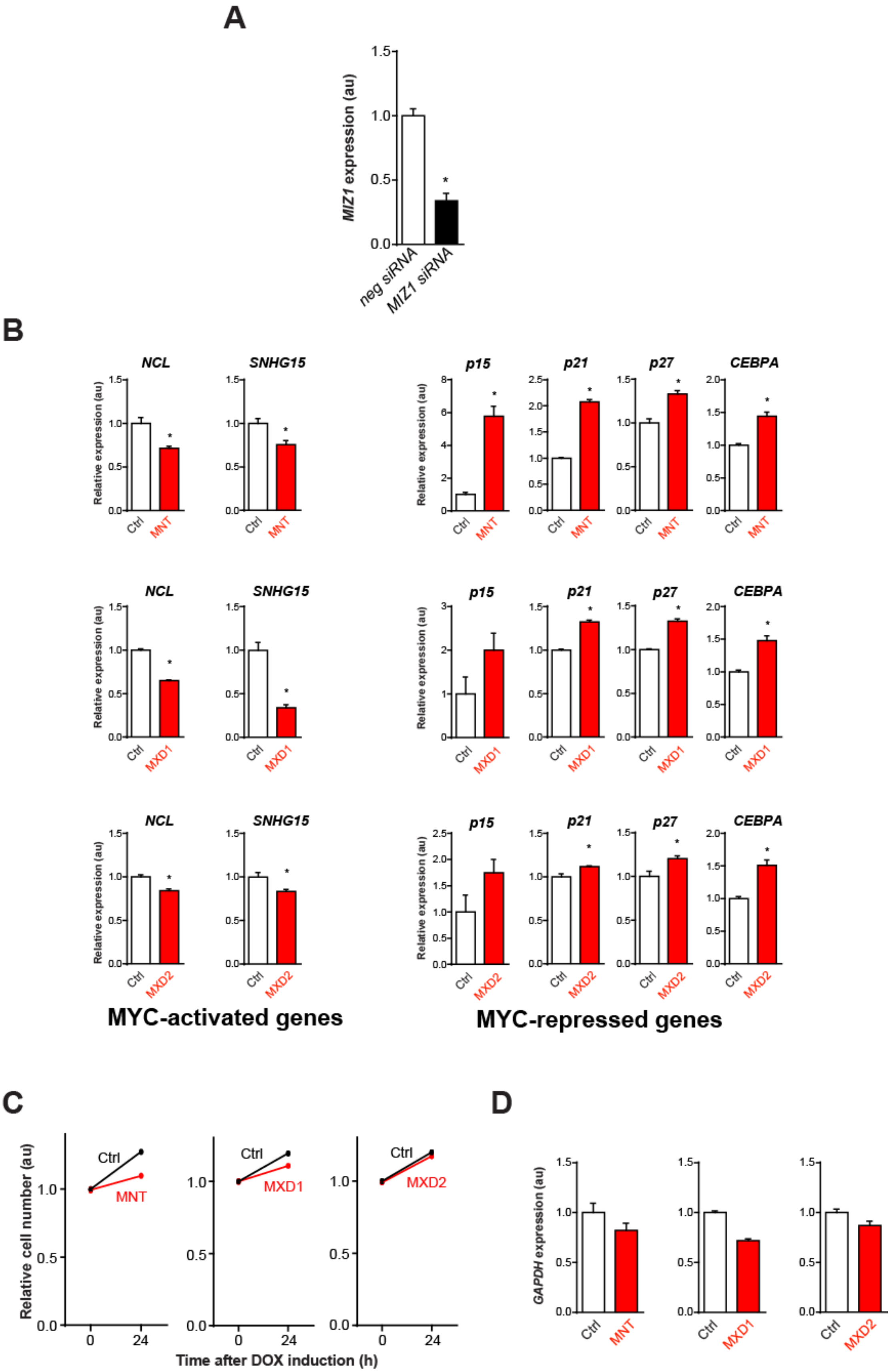

**Supplemental Figure S2. (A)** qPCR quantification of *MIZ1* mRNA in U2OStx cells transfected with siRNA against *MIZ1* (n=3). **(B)** Expression of MYC-activated and MYC-repressed genes (24h) determined by qPCR in U2OStx with DOX-induced MXDs (n=3). **(C)** Growth rate of U2OStx cells expressing MNT, MXD1, and MXD2 determined by WST-8 assay. Cells were treated with DOX or PBS (Ctrl) for 24 h. **(D)** *GAPDH* mRNA abundance in U2OStx cells 24 hours after induction of MXDs. Data are presented as mean ± SEM. * $P < 0.05$; Student's *t*-test.

Fig. S3 Shostak et al

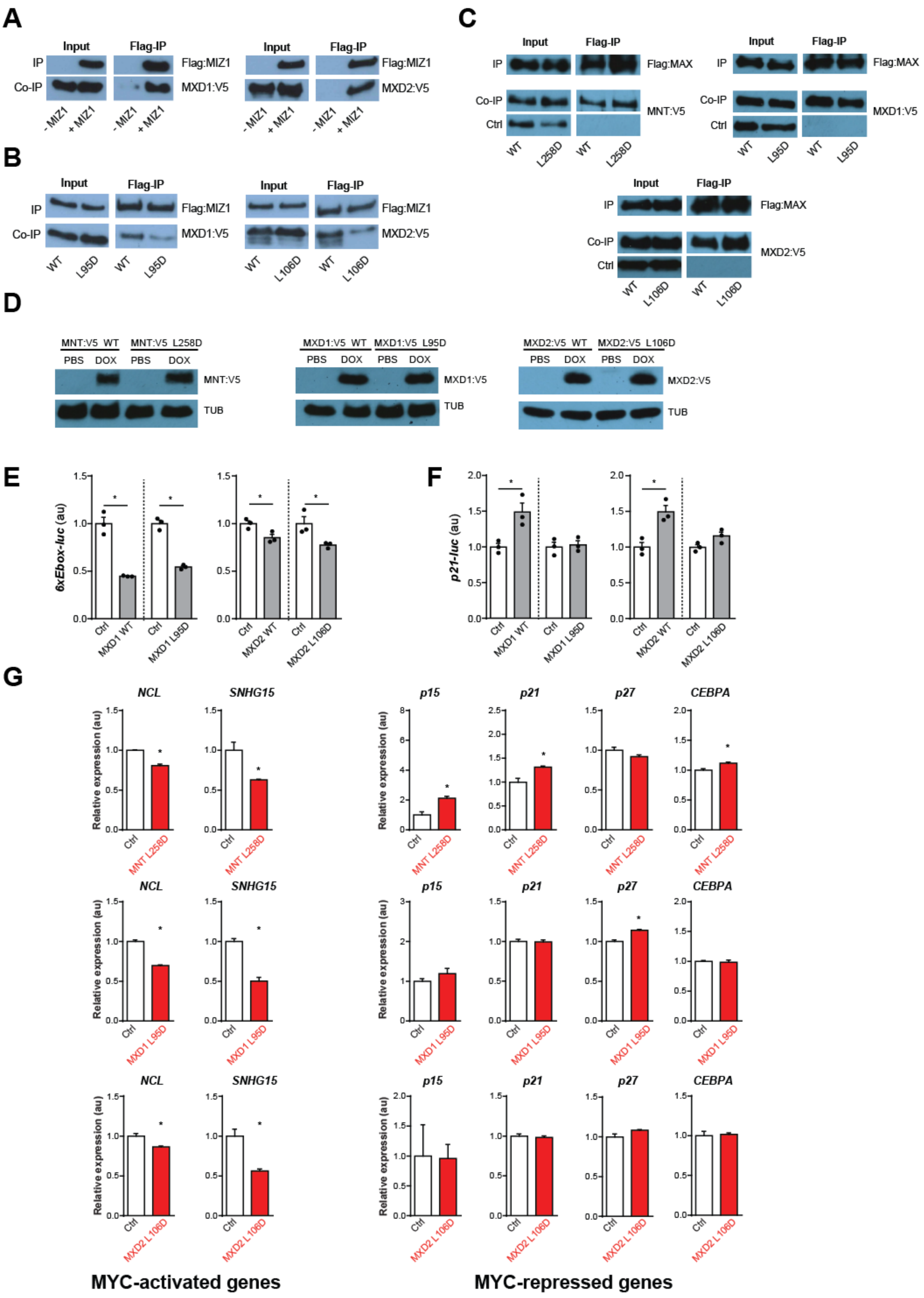

**Supplemental Figure S3. (A)** Co-immunoprecipitation of V5-tagged MXD1 and MXD2 with FLAG-tagged MIZ1 from HEK293 lysates. **(B)** FLAG:MIZ1 pulldown and co-immunoprecipitation of WT and L-D versions of MXD1 and MXD2 expressed in HEK293 cells. MIZ1 and MXDs were tagged with FLAG and V5 epitopes, respectively. **(C)** Co-immunoprecipitation of WT and L-D versions of MNT, MXD1, and MXD2 with MAX expressed in HEK293 cells. MXDs and MAX were tagged with V5 and FLAG epitopes, respectively. Ctrl: Flag-IPs were performed from cells transfected with only V5-taged MXDs. cells. **(D)** WT and L-D mutants of MNT, MXD1, and MXD2 are expressed at similar levels. Western blot analysis of U2OStx cell lysates expressing DOX-induced WT or L-D versions of MXDs. Repression of *6xEbox-luc* **(E)** and activation of *p21-luc* **(F)** in U2OS cells overexpressing WT or L-D mutants of MXD1 and MXD2 after induction with DOX after 24 hours (n=3). **(G)** qPCR quantification of MYC-activated and MYC-repressed genes (24 h) in U2OStx expressing L-D versions of MXDs (n=3). Data are presented as mean ± SEM. * $P < 0.05$; one-way ANOVA with Bonferroni post-test (**E** and **F**) and Student's *t*-test (**G**).

Fig. S4 Shostak et al

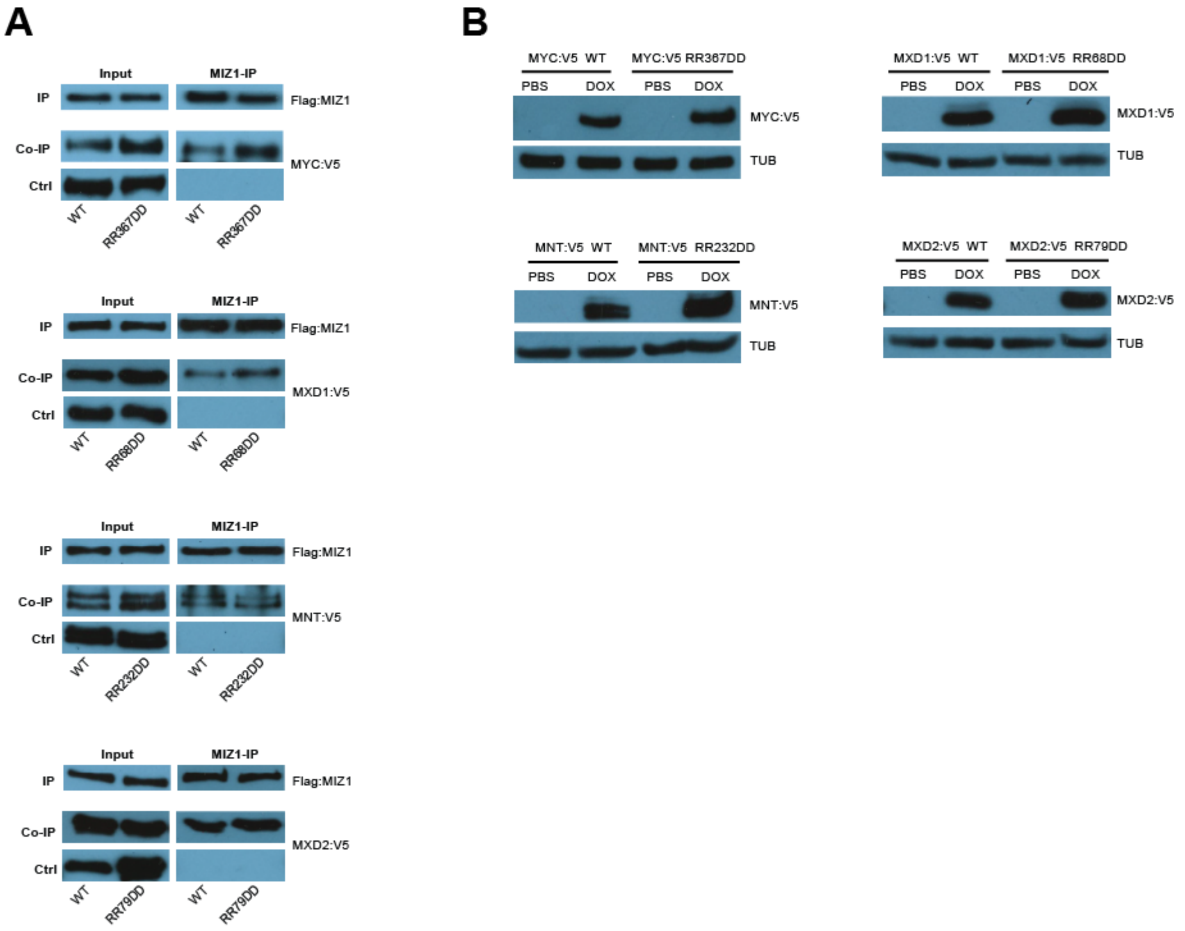


**Supplemental Figure S4. (A)** Co-immunoprecipitation of WT and RRDD versions of V5 tagged MYC, MNT, MXD1, and MXD2 with FLAG:MIZ1 expressed in HEK293 cells. Ctrl: Flag-IPs were performed from cells expressing only indicated V5-tagged MXDs. **(B)** Western blot analysis of DOX-induced MYC, MNT, MXD1, and MXD2 in U2OStx cells. Tubulin (TUB) is shown for control.

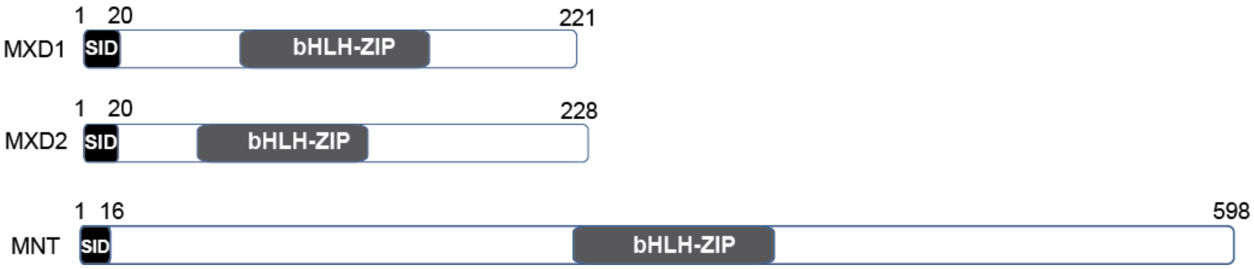


**Supplemental Figure S5.** Schematic of the domain structure of MXDs. MXDs contain a central bHLH-Zip DNA-binding domain (bHLH) and an N-terminal SID domain. SID domains are short segments of ~20 aa required for the recruitment of mSIN3-HDAC co-repressor complexes.

Fig. S6 Shostak et al

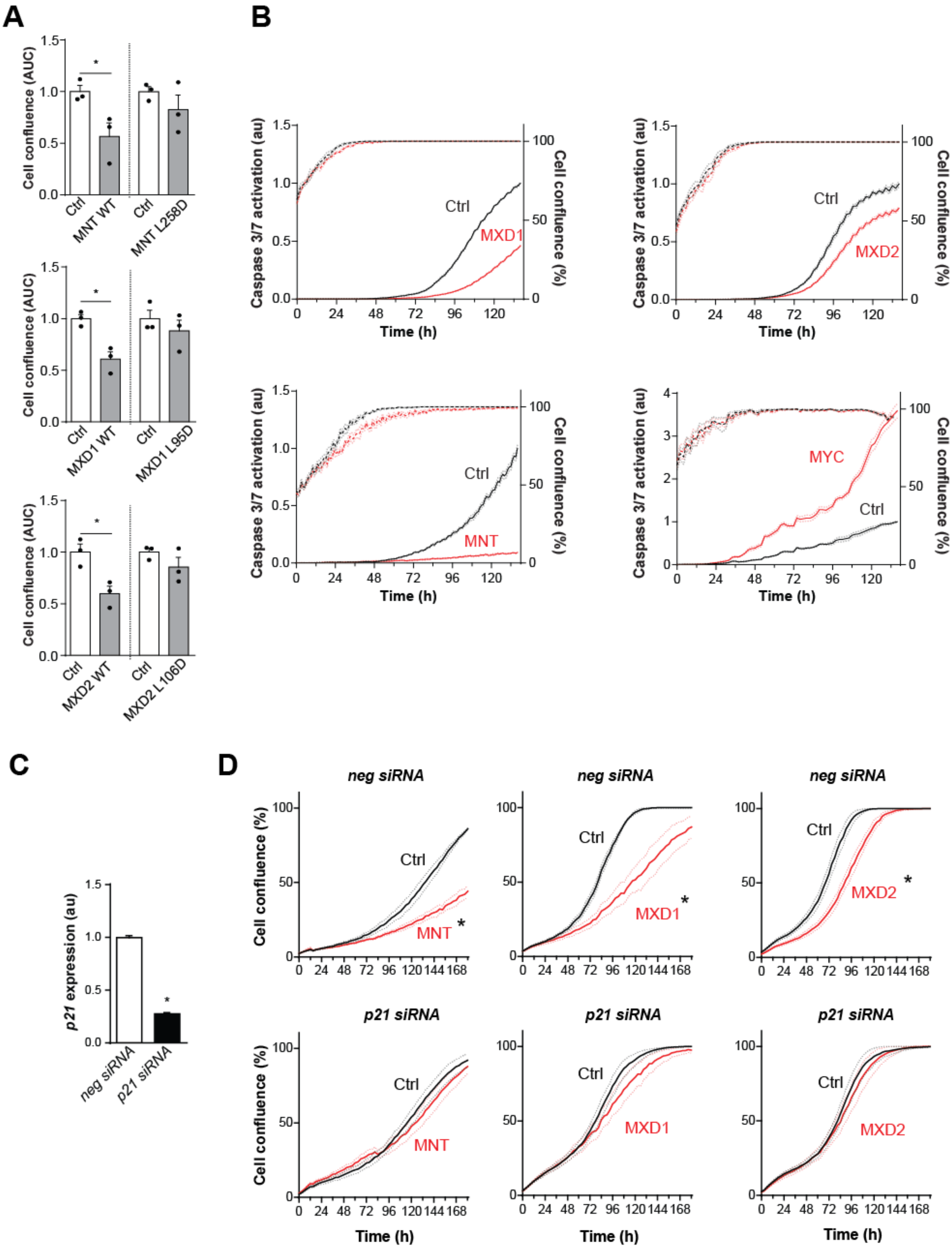

**Supplemental Figure S6. (A)** L-D versions of MXDs do not inhibit proliferation. Relative confluence (AUC) of U2OStx cells expressing DOX-induced WT and L-D versions MXDs over several days is shown (n=3). For raw data see Fig. 6A. **(B)** Normalized Caspase 3/7 activity (solid lines) and confluence (dashed lines) in U2OStx cells expressing inducible MNT, MXD1, MXD2, and MYC. Cells, seeded at 50-60% confluency one day before the experiment, were treated with DOX or PBS (Ctrl) at the time point 0 and green fluorescence was measured using the Incucyte ZOOM system (n=3). **(C)** qPCR analysis showing downregulation of *p21* by siRNA in U2OStx cells (n=3). * $P < 0.05$; Student's *t*-test. **(D)** Inhibition of cell proliferation by MIZ1 and MXDs requires p21. Growth curves of indicated transgenic cell lines treated with siRNA against *p21* (n=3). * $P < 0.05$; AUC comparison of DOX vs. Ctrl curves by two-way ANOVA with Bonferroni post-test. Data are presented as mean ± SEM.

### Supplemental Table 1. Primer sequences and siRNAs.

| qPCR primers for gene expression analysis | | |
|---|---|---|
| **name** | **sequence** | **source** |
| *GAPDH_F* | TGCACCACCAACTGCTTAGC | Zhang, E.E. *et al.* (2009) A genome-wide RNAi screen for modifiers of the circadian clock in human cells. *Cell* |
| *GAPDH_R* | ACAGTCTTCTGGGTGGCAGTG | |
| *p15_F* | CTAGTGGAGAAGGTGCGACAGC | Zhang, W. *et al.* (2012) Variants on Chromosome 9p21.3 Correlated With ANRIL Expression Contribute to Stroke Risk and Recurrence in a Large Prospective Stroke Population. *Stroke* |
| *p15_R* | CACCAGCGTGTCCAGGAAG | |
| *p27_F* | GGTTAGCGGAGCAATGCG | Khattar, E. *et al.* (2010) Mitogenic regulation of p27(Kip1) gene is mediated by AP-1 transcription factors. *J Biol Chem* |
| *p27_R* | TCCACAGAACCGGCATTTG | |
| *p21_F* | TGGAGACTCTCAGGGTCGAAA | Al-Haj, L. et al. (2012) Regulation of p21/CIP1/WAF-1 mediated cell-cycle arrest by RNase L and tristetraprolin, and involvement of AU-rich elements. *Nucleic Acids Res* |
| *P21 _R* | GGCGTTTGGAGTGGTAGAAATC | |
| *MIZ1_F* | TGAAGATCCACATCGCTGACG | Walz, S. et al. (2014) Activation and repression by oncogenic MYC shape tumour-specific gene expression profiles. *Nature* |
| *MIZ1_R* | GGTCTGCAAACTGTCGCTG | |
| *SNHG15_F* | CTCCGTACTCCGTACTTCGT | This work |
| *SNHG15_R* | GGGGTGTTCAGCAACTATTC | |
| *NCL_F* | GCACCTGGAAAACGAAAGAAGG | PrimerBank ID55956787c2 |
| *NCL_R* | GAAAGCCGTAGTCGGTTCTGT | |
| *CEBPA_F* | AACATCGCGGTGCGCAAGAG | Placke, T. et al. (2014) Requirement for CDK6 in MLL-rearranged acute myeloid leukemia. *Blood* |
| *CEBPA_R* | TTCGCGGCTCAGCTGTTCCA | |
| *MNT_F* | TCGGAACCAGAGAAGTCCAC | Menssen, A. and Hermeking, H. (2002) Characterization of the c-MYC-regulated transcriptome by SAGE: Identification and analysis of c-MYC target genes. *PNAS* |
| *MNT_R* | CGCTCCATTTCATGCTCATA | |
| *MXD1_F* | ACATGGTTATGCCTCCATGTTAC | Wang, X. et al. (2013) Hypermethylation reduces expression of tumor-suppressor PLZF and regulates proliferation and apoptosis in non-small-cell lung cancers. *FASEB J* |
| *MXD1_R* | AGATGAGCCCGTCTATTCTTCTC | |
| *MXD2_F* | ATTCCACTAGGACCAGACTGC | Tsao, CC. et al. (2008) Inhibition of Mxi1 suppresses HIF-2alpha-dependent renal cancer tumorigenesis. *Cancer Biol Ther* |
| *MXD2_R* | CTGGTGGTACTTATATTGTCCAC | |

| qPCR primers for ChIP | | |
|---|---|---|
| **name** | **sequence** | |
| *p21_TSS_MIZ1bs_F* | CCGAAGTCAGTTCCTTGTGG | Walz, S. *et al.* (2014) Activation and repression by oncogenic MYC shape tumour-specific gene expression profiles. *Nature* |
| *p21_TSS_MIZ1bs_F* | CGCTCTCTCACCTCCTCTGA | |
| *neg region chr11_F* | TTTTCTCACATTGCCCCTGT | |
| *neg region chr11_R* | TCAATGCTGTACCAGGCAAA | |
| *NCL_Ebox_F* | GGGACTCGACTCCTGACG | Shostak, A. *et al.* (2016) MYC/MIZ1-dependent gene repression inversely coordinates the circadian clock with cell cycle and proliferation. *Nature communications* |
| *NCL_Ebox_R* | ACTCCGACTAGGGCCGATAC | |

| siRNA sequences | | | |
|---|---|---|---|
| *negative siRNA* | N/A | 20 µM | Silencer Select Negative Control No. 1 siRNA, cat # 4390843, Ambion |
| *MYC* | N/A | 10 µM | sc-29226, SantaCruz |
| *MIZ1 pool* | AGUUCAACCAGGUAGGGAAdtdt<br>GGUGGACGGUGUUCACUUUdtdt | 25 + 25 µM | Kaur, M. *et al.* (2013) MYC acts via the PTEN tumor suppressor to elicit autoregulation and genome-wide gene repression by activation of the Ezh2 methyltransferase. *Cancer Research.*<br>Walz, S. *et al.* (2014) Activation and repression by oncogenic MYC shape tumour-specific gene expression profiles. *Nature* |
| *p21(CDKN1A)* | AACAUACUGGCCUGGACUGUUdtdt | 50 µM | Wall, SJ. *et al.* (2007) The cyclin-dependent kinase inhibitors p15INK4B and p21CIP1 are critical regulators of fibrillar collagen-induced tumor cell cycle arrest. *The Journal of Biological Chemistry* |
| *MNT* | CCUCGGAAAUCAGUGCGAUdtdt | 50 µM | Wu, J. *et al.* (2012) MNT inhibits the migration of human hepatocellular carcinoma SMMC7721 cells. *Biochemical and Biophysical Research Communications* |
| *MXD1* | GAGAGAAGCUGAACAUGGUUdtdt | 50 µM | Xu, L. *et al.* (2007) c-IAP1 Cooperates with Myc by Acting as a Ubiquitin Ligase for Mad1. *Molecular Cell* |
| *MXD2 (MXI1)* | GGAGAUGGAACGAAUACGAdtdt | 50 µM | Corn, PG. *et al.* (2005). Mxi1 is Induced by Hypoxia in a HIF-1-Dependent Manner and Protects Cells from c-Myc-Induced Apoptosis. *Cancer Biology & Therapy* |
